# Archaeal community structure and underlying ecological processes in swine manure slurry

**DOI:** 10.1101/233445

**Authors:** Priyanka Kumari, Hong L. Choi

## Abstract

The ecological processes underlying the observed patterns in community composition of archaea in swine manure slurry are poorly understood. We studied the archaeal communities from six swine manure slurry storage tanks using paired-end Illumina sequencing of the v3 hypervariable region of 16S rRNA gene. Across all samples, the archaeal community was dominated by methanogens related to *Thermoplasmata, Methanomicrobia*, and *Methanobacteria* classes. At the genus level, the archaeal community was dominated by a single uncultured lineage of archaea, *vadinCA11*, followed by methanogenic genera *Methanobrevibacter*, *Methanosarcina*, *Methanosphaera*, *Methanogenium*, *Methanocorpusculum*, *Methanoculleus*, and *Methanomicrococcus*. Significant phylogenetic signals were detected across relatively short phylogenetic distances, indicating that closely related archaeal operational taxonomic units (OTUs) tend to have similar niches. The standardized effect sizes of mean nearest taxon distance (SES.MNTD) showed that archaeal community was phylogenetically clustered, suggesting that environmental filtering deterministically influence the within-community composition of archaea. However, between-community analysis based on *β*-nearest taxon index (*β*NTI) revealed that both deterministic selection and stochastic processes operate simultaneously to govern the assembly of archaeal communities. These findings significantly enhance our understanding of archaeal community assembly and underlying ecological processes is swine manure slurry.

## Introduction

Archaea constitute a small fraction of the microbial biomass on Earth, however they involved in various essential ecological processes including nitrification, denitrification and methanogenesis (Offre et al. 2013). Originally, it was believed that archaea inhabit only under extreme conditions, however, now they are known to be present in wide variety of ecosystems including soil (Bintrim et al. 1997; Bates et al. 2011; Tripathi et al. 2013), marine and fresh-water habitats (Lipp et al. 2008; Lliros et al. 2008; Teske and S0rensen 2008; Auguet et al. 2011), animal feces and stored manure (Mao et al. 2011; Barret et al. 2015), and intestinal tracts of animals (Shin et al. 2004; Pei et al. 2010). Culture-independent high-throughput sequencing methods have significantly improved our understanding of the diversity and composition of archaea across various ecosystems (Auguet et al. 2010). However, comparatively little is known about the ecological processes that shape the assembly of archaeal communities.

The assembly of microbial communities is influenced by two types of ecological processes-deterministic selection and stochastic processes (Stegen et al. 2012; Wang et al. 2013). Traditionally, the assembly of microbial communities has been thought to be influenced primarily by deterministic selection (Baas-Becking 1934; Torsvik et al. 2002), in which certain abiotic and biotic factors select a particular assemblage of microbial species (Fierer and Jackson 2006; Lozupone and Knight 2007). However, some studies have reported that stochastic processes (e.g., random birth, death, dispersal, and colonization) can also play a major role in shaping assembly of microbial communities (Peay et al. 2010; Caruso et al. 2011). More recently, it has been found that both deterministic and stochastic processes can act concurrently to influence the community assembly of microbial communities (Chase and Myers 2011; Langenheder and Székely 2011; Stegen et al. 2012; Zhou et al. 2014).

Modern swine production facilities are more intensified and generate large quantities of manure slurry (Barker and Overcash 2007). The swine manure slurry is commonly stored in deep pits for months before being applied to agricultural lands as fertilizer. During storage the swine manure slurry decomposes, and emits a considerable quantity of methane (EPA 2013), which is exclusively generated by methanogenic archaea (Whitman et al. 2006). Over the past decade, several studies have characterized the archaeal community present in swine feces and stored manure slurry by using both cultivation-based (Miller et al. 1986) and culture-independent methods (Whitehead and Cotta 1999; Mao et al. 2011; Da Silva et al. 2014). However, it is still not clear what underlying ecological processes are important in structuring the community composition of archaea in swine manure slurry.

Therefore, this study was conducted to answer the following questions:

1. What are the dominant archaeal taxa present in swine manure slurry?
2. How does the community composition of archaea is governed by the underlying ecological processes (deterministic vs. stochastic) in both within- and between-community-level analyses?

We applied an ecological null modeling approach to understand the relative influence of deterministic selection and stochastic processes in structuring the archaeal communities (Webb et al. 2002; Stegen et al. 2012).

## Material and Methods

### Sample collection

Swine manure slurry samples were collected from six different commercial pig farms in South Korea in June 2013. For each sample, slurry was collected from the top 1 m of the storage tank at five different points, and finally one liter of slurry was collected after through mixing (for detailed information on sampling, see (Kumari et al. 2015). After collection the swine manure slurry samples were kept in an ice box and transported to the laboratory for DNA extraction.

### DNA extraction, PCR amplification, and Illumina sequencing

DNA was extracted from the pellet of centrifuged (at 14,000 x g for 5 min) swine manure slurry samples (5-ml each) using a PowerSoil DNA isolation kit (MoBio Laboratories, USA). The v3 hypervariable region of archaeal 16S rRNA gene was amplified using primer pair S-D-Arch-0349-a-S-17 (5′GYGCASCAGKCGMGAAW3′)/ S-D-Arch-0519-a-A-16 (5′TTACCGCGGCKGCTG3′) (Klindworth et al. 2012). The resulting amplicons were sequenced at the Beijing Genome Institute (BGI) (Hong Kong, China) using Hiseq™ 2500 platform (Illumina, USA) to generate 150 bp paired-end reads. The raw sequence data were deposited into the NCBI short reads archive (SRA) database under accession number SRP117664.

### Sequence processing

The paired-end sequences were assembled using PANDAseq (Masella et al. 2012), and subsequently processed in Mothur (Schloss et al. 2009). The assembled 16S rRNA gene sequences were aligned against SILVA reference alignment v123 (http://www.arb-silva.de/). The alignment was screened for chimeric sequences using ‘chimera.uchime’ command implemented in mothur in de novo mode (Edgar et al. 2011). The resulting alignment was classified against Greengenes reference taxonomy database (release gg_13_8_99; August 2013). The sequences were grouped into operational taxonomic units (OTUs) at ≥97% sequence similarity using the average neighbor clustering algorithm. Singleton OTUs were removed to avoid spurious results due to sequencing errors. Finally, the dataset was rarified to a depth of 10,291 sequences per sample using the ‘sub.sample’ command in mothur.

### Phylogenetic and statistical analysis

A maximum-likelihood tree was constructed from the sequence alignment of representative OTUs using FastTree (Price et al. 2010). For evaluating phylogenetic signals, environmental-optima for all OTUs were calculated by following the procedure described by Stegen et al. (2012). Then, among-OTU differences in environmental optima were calculated as Euclidean distances. Mantel correlogram was used to assess the correlation coefficients between differences in environmental optima and phylogenetic distances with 999 permutations. We calculated mean nearest taxon distance (MNTD) to emphasize phylogenetic relationships between archaeal OTUs (Webb et al. 2002). To analyze the turnover in phylogenetic community composition, *β*MNTD was calculated using ‘comdistnt’ function (abundance.weighted = TRUE) in ‘picante’ R package (Kembel et al. 2010). Whereas, Bray-Curtis distance was calculated using ‘vegdist’ function in ‘vegan’ R package (Oksanen et al. 2013) to evaluate the turnover in taxonomic community composition (at the OTU level). Nonmetric multidimensional scaling (NMDS) plots were used to visualize the taxonomic and phylogenetic composition of archaeal community across all samples. To assess congruence among taxonomic and phylogenetic NMDS ordinations, we used ‘procrustes’ function in vegan R package.

The standardized effect sizes of MNTD (SES.MNTD) was calculated using the null model ‘taxa.labels’ (999 randomization) in ‘picante’ R package to evaluate the phylogenetic community assembly. The mean value of SES.MNTD across all communities deviating significantly from zero indicates clustering (SES.MNTD<0) or over dispersion (SES.MNTD>0) (Webb et al. 2002; Kembel et al. 2010).

Furthermore, a null modeling approach was employed to disentangle the community assembly processes using R software (Stegen et al. 2012; Stegen et al. 2013; Wang et al. 2013). To infer this, the *β*-nearest taxon index (*β*NTI) was calculated which estimates the deviation of the observed *β*MNTD from the null distribution of *β*MNTD (using 999 randomizations). The |*β*NTI| values > 2 indicate deterministic selection (*β*NTI <-2: homogeneous selection; *β*NTI >+2: variable selection) is responsible for differences in community composition, whereas |*β*NTI| values <2 indicate that the observed difference in community composition is result of stochastic processes (Dini-Andreote et al. 2015).

## Results and discussion

A total of 122,321 archaeal 16S rRNA gene sequences (with an average length of 191 bp) were obtained from six swine manure slurry samples, with coverage ranging from 10,896 to 26,661 reads per sample. Despite the high level of sequencing coverage, the rarefaction curves did not reach a plateau (Fig. 1), indicating that the actual archaeal diversity is much higher. Of the 122,321 sequences, around 98.8% sequences were classified up to phylum *Euryarchaeota. Euryarchaeota* are known to inhabit pig feces and mostly dominated by the methanogenic members of this phylum (Cardinali-Rezende et al. 2012; Da Silva et al. 2014).

**Fig. 1.**
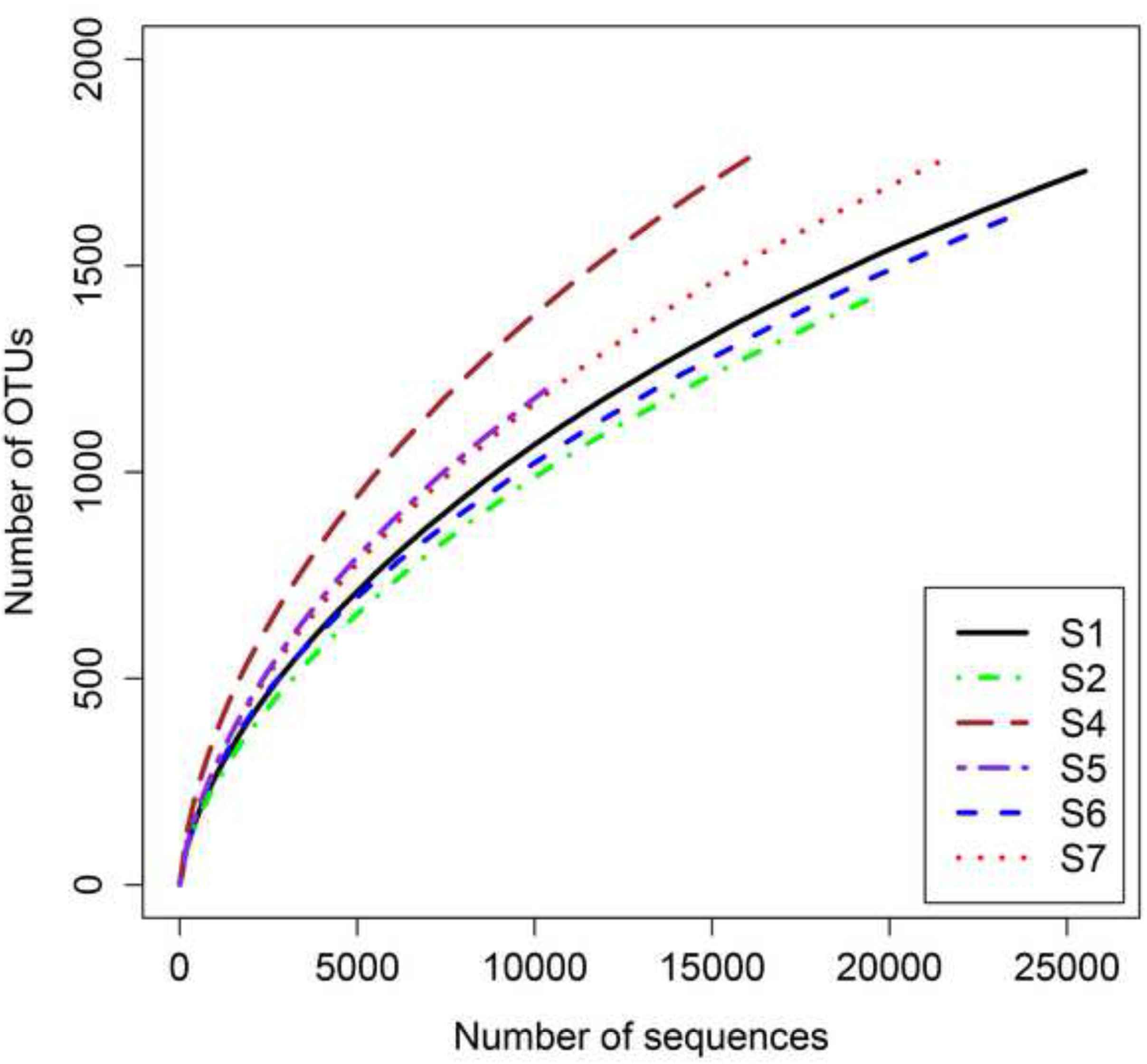
Rarefaction curves of archaeal operational taxonomic units (≥97% sequence similarity) for each sample of swine manure slurry.

Across all samples, *Thermoplasmata* was the most abundant euryarchaeal class (75,127 sequences, 61.4% of all sequences) followed by *Methanomicrobia* (22,432 sequences, 18.3% of all sequences) and *Methanobacteria* (16,226 sequences, 13.3% of all sequences) (Fig. 2). All these archaeal classes have been reported to dominate pig feces (Dridi et al. 2009; Mao et al. 2011). The candidate genus *vadinCA11* (Godon et al. 1997) predominated across all samples representing around 56% of the whole archaeal community (Fig.3). The dominance of vadinCA11 is previously documented in swine manure and storage pits (Whitehead and Cotta 1999; Snell-Castro et al. 2005). So far, the metabolic activity of this uncultured archaeal lineage is remains unclear. The other most dominant archaeal genera detected across all samples were *Methanobrevibacter, Methanosarcina, Methanosphaera, Methanogenium, Methanocorpusculum, Methanoculleus*, and *Methanomicrococcus* (Fig.3). All these genera use hydrogenotrophic methanogenesis pathway except *Methanosarcina*, which can use both the acetoclastic and the hydrogenotrophic methanogenesis pathways (Phelps et al. 1985).

**Fig. 2.**
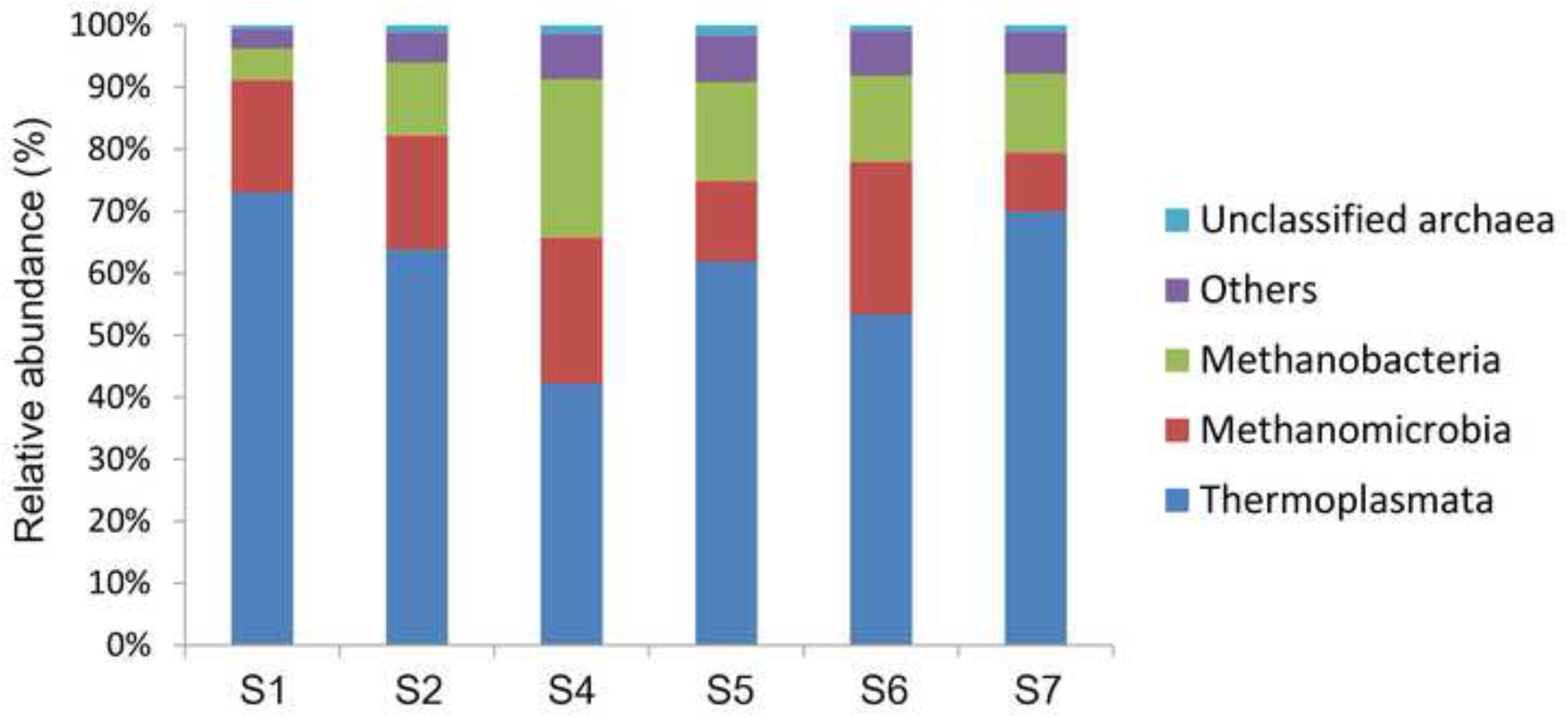
The relative abundance of dominant archaeal taxa (at class level) present in each sample of swine manure slurry.

**Fig. 3.**
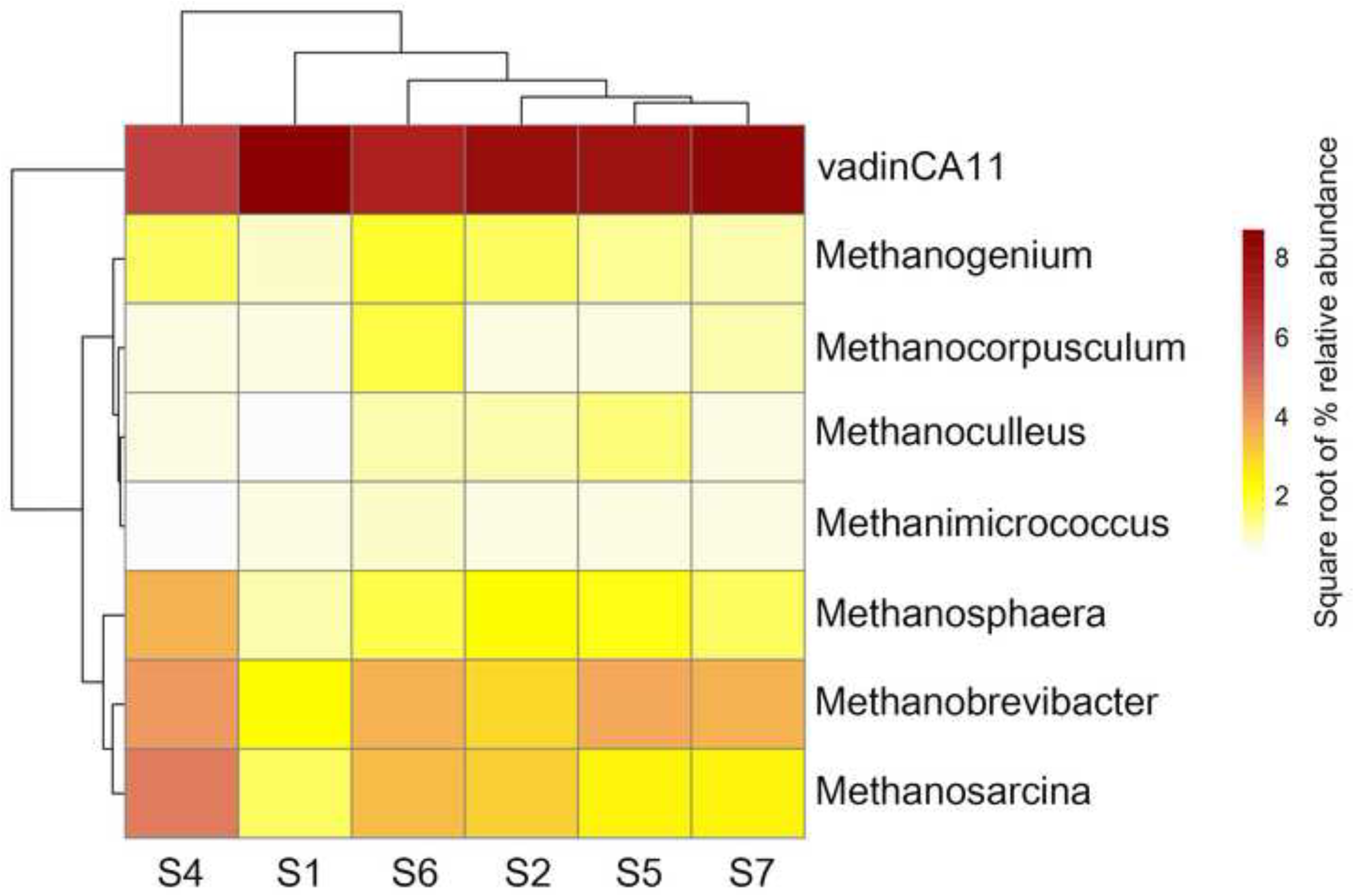
Heat map showing the relative abundance (square root transformed) of the most dominant archaeal genera in each sample of swine manure slurry.

Testing phylogenetic signal in the ecological niches of species is a prerequisite for making ecological inferences using phylogenetic information (Cavender-Bares et al. 2009; Fine and Kembel 2011). Mantel correlogram showed significant phylogenetic signal across relatively short phylogenetic distances (Fig.4), which indicates that niche preferences of closely related archaeal OTUs are more similar to each other than to the niche preferences of distant relatives (Losos 2008). Therefore we calculated MNTD to quantify the phylogenetic distances among close relatives (Webb et al. 2002). This result is consistent with the observation of our previous study on bacterial communities in swine slurries (Kumari et al. 2015), which also found significant phylogenetic signal across relatively short phylogenetic distances. Furthermore, similar results were also reported on archaeal communities in soil ecosystems (Tripathi et al. 2015).

**Fig. 4.**
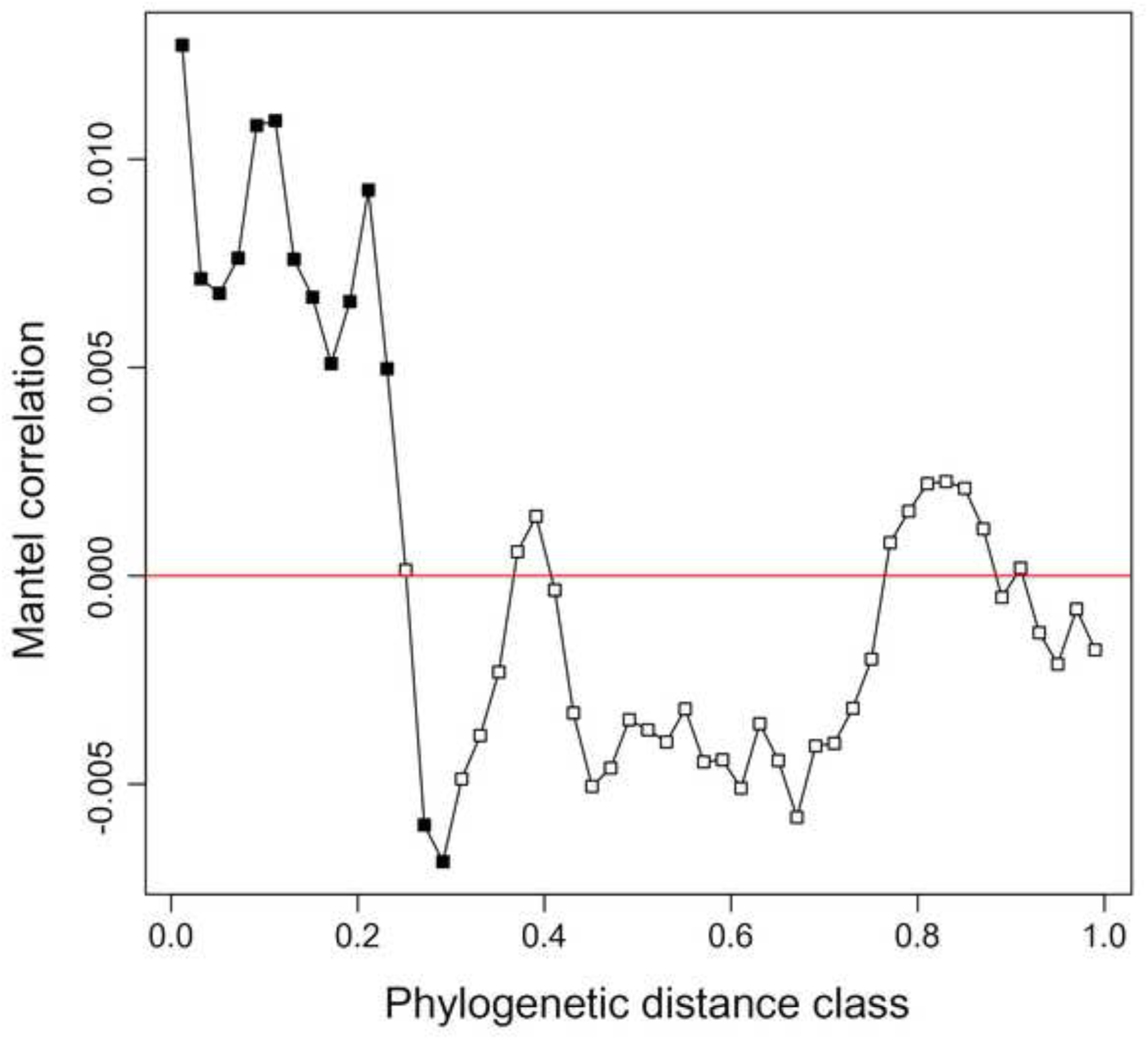
Mantel correlogram between differences in environmental optima of OTUs and phylogenetic distances of archaea in swine manure slurry. Closed squares represent significant (*P* < 0.05) phylogenetic signals.

Procrustes analysis comparing spatial fit between NMDS plots generated using Bray-Curtis (taxonomic) and *β*MNTD (phylogenetic) distances showed concordance (procrustes correlation, *R* = 0.80, *P* = 0.04; Fig. 5), indicating that archaeal communities in swine slurry samples have a strong evolutionary structural component. The SES.MNTD distribution mean was significantly less than zero across all samples (Fig. 6a; one-tailed t-test, *P* < 0.05), indicating that the each archaeal community was phylogenetically clustered largely due to environmental filtering. Environmental filtering play a key role in shaping species community assembly (Webb et al. 2002).This result is consistent with previous studies on microbial communities in different environments (Kumari et al. 2015; Ren et al. 2015; Yan et al. 2016), which showed that microbial communities had a tendency to be phylogenetically clustered.

**Fig. 5.**
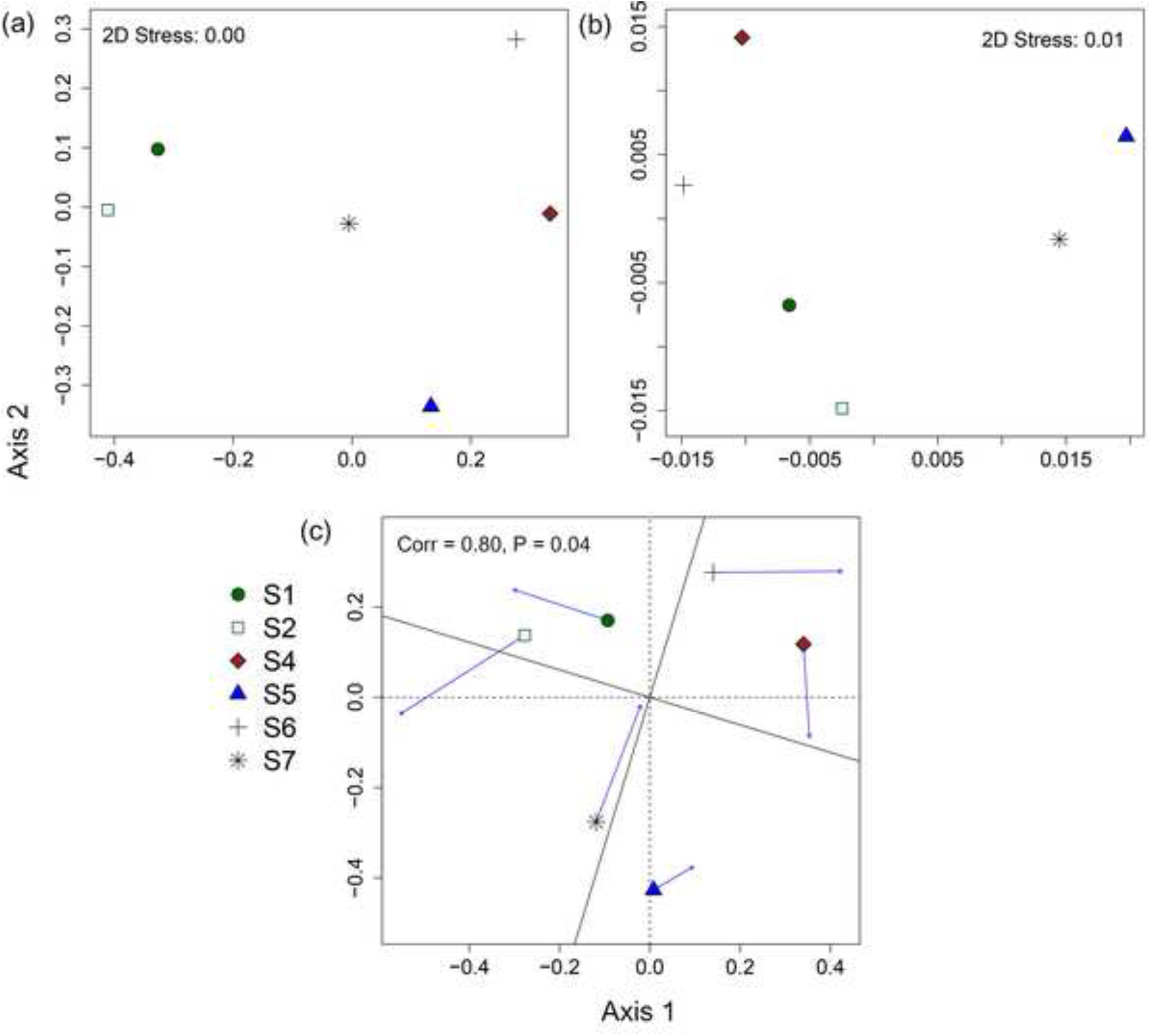
NMDS ordinations showing the first two axes of (a) taxonomic community composition based on Bray-Curtis distance; (**b**) phylogenetic community composition based on *β*MNTD and (c) Procrustes analysis comparing taxonomic and phylogentic community composition of archaea. The arrows in Procrustes analysis point towards the target configuration (taxonomic community composition), and symbols represent the rotated configuration (phylogenetic community composition). Correlation and significance values were calculated using the ‘protest’ function in vegan R package.

**Fig. 6.**
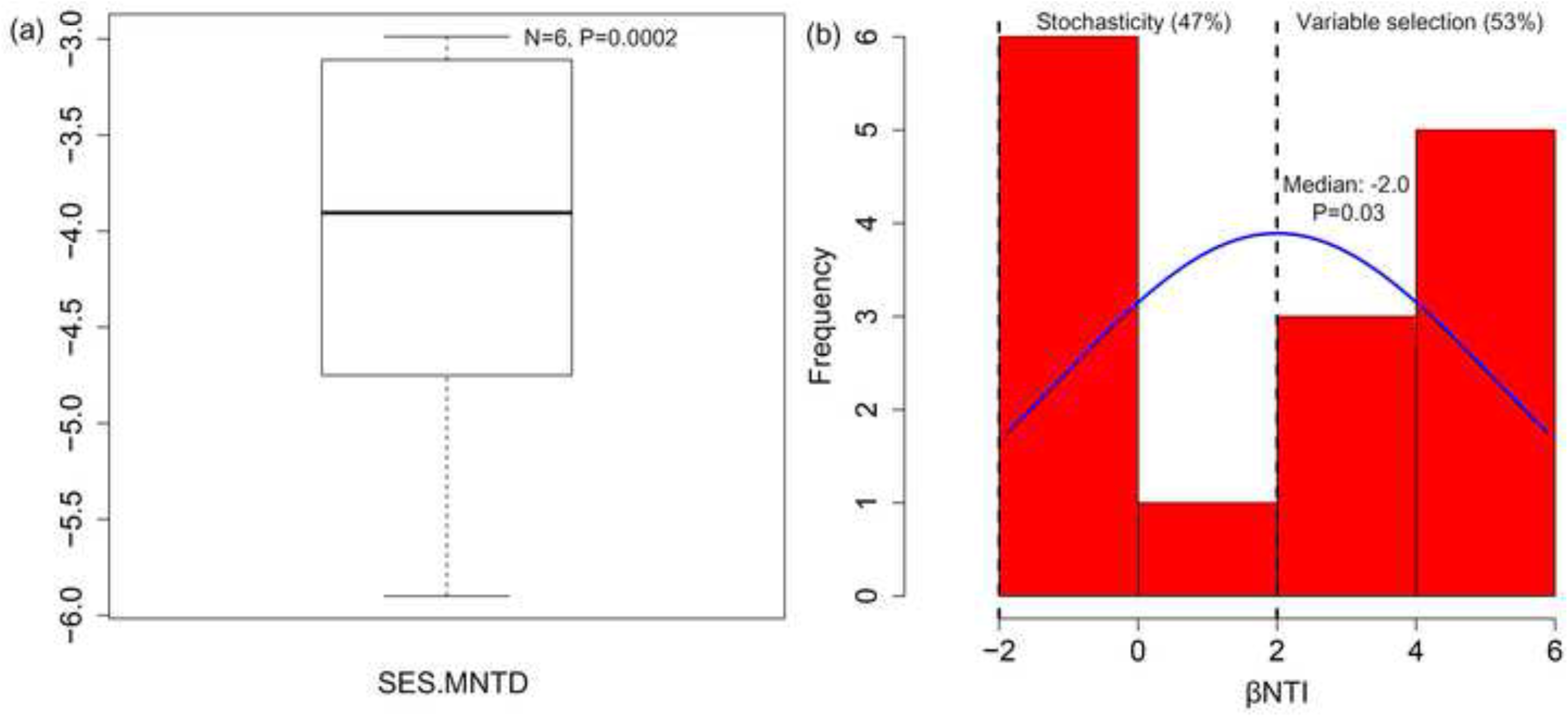
(a) Box plot showing variation in SES.MNTD values across all samples. The SES.MNTD distribution mean was significantly less than zero (one-tailed t-test, *P* < 0.05). (b) Frequency estimates for distributions of *β*NTI. The median of the *β*NTI distribution was significantly different from expected value of zero (one-tailed t-test, *P* < 0.0001).

To infer the relative influence of deterministic selection vs. stochastic processes that govern the phylogenetic turnover of archaeal communities in swine manure slurry, we employed between-community null modeling approach and calculated *β*NTI (Stegen et al. 2012; Stegen et al. 2013; Wang et al. 2013). The median of the *β*NTI distribution was significantly greater than the expected value of zero (median *βNTI*= +2.00, one-tailed t-test, p< 0.0001; Fig. 6b), indicating that on average variable selection governs archaeal community dynamics. The physicochemical properties of swine manure slurry varied widely across samples (Suresh and Choi 2011), and this heterogeneous condition among swine manure slurry samples could results in strong influence of variable selection. We also quantified the relative proportions of both ecological processes across all comparisons (Fig. 6b). The results showed that though the relative contribution of deterministic selection was higher (variable selection = 53%), stochastic processes (47%) also appeared to influence the turnover of archaeal communities. These results are consistent with the recent theoretical frameworks that both deterministic selection and stochastic processes govern the assembly of microbial communities (Chase and Myers 2011; Langenheder and Székely 2011; Stegen et al. 2012; Zhou et al. 2014).

## Conclusions

In conclusion, we found that the archaeal community in swine manure slurry was dominated by methanogenic archaea. The phylogenetic signals detected across relatively short phylogenetic distances revealed that closely related archaeal OTUs have more similar niche preferences, hence they are ecologically similar. The archaeal community was phylogenetically clustered within-community analysis (SES.MNTD) and thus environmental filtering deterministically governs the assembly of each community in swine manure slurry. However, relative to within-community analysis, the turnover between archaeal communities (*β*NTI) was governed by both deterministic selection and stochastic processes. Overall, this study provides valuable insights into the community assembly and ecology of archaea in swine manure slurry.

## Acknowledgements

This work was supported by Korea Institute of Planning and Evaluation for Technology in Food, Agriculture, Forestry, and Fisheries (IPET) from the Ministry of Agriculture, Food, and Rural Affairs (MAFRA) through project no. 312036-03-2-HD030.

## References

Auguet JC, Nomokonova N, Camarero L, Casamayor EO (2011) Seasonal changes of freshwater ammonia-oxidizing archaeal assemblages and nitrogen species in oligotrophic alpine lakes. Appl Environ Microbiol 77:1937–1945

Auguet JC, Barberan A, Casamayor EO (2010) Global ecological patterns in uncultured Archaea. ISME J 4:182–190

Baas-Becking LGM (1934) Geobiologie of inleiding tot de milieukunde. Van Stockum WP & Zoon NV, The Hague

Barker J, Overcash M (2007) Swine waste characterization: a review. T ASABE 50:651–657

Barret M, Gagnona N, Morissettea B, Kalmokoffb ML, Stephen ET, Brooks PJ et al. (2015) Phylogenetic identification of methanogens assimilating acetate-derived carbon in dairy and swine manures. Syst Appl Microbiol 38:56–66

Bates ST, Berg-Lyons D, Caporaso JG, Walters WA, Knight R, Fierer N (2011) Examining the global distribution of dominant archaeal populations in soil. ISME J 5:908–917

Bintrim SB, Donohue TJ, Handelsman J, Roberts GP, Goodman RM (1997) Molecular phylogeny of Archaea from soil. Proc Natl Acad Sci USA 94:277–282

Cardinali-Rezende J, Pereira ZL, Sanz JL, Chartone-Souza E, Nascimento AM (2012) Bacterial and archaeal phylogenetic diversity associated with swine sludge from an anaerobic treatment lagoon. World J Microbiol Biotechnol 28:3187–3195

Caruso T, Chan Y, Lacap DC, Lau MC, McKay CP, Pointing SB (2011) Stochastic and deterministic processes interact in the assembly of desert microbial communities on a global scale. ISME J 5:1406–1413

Cavender-Bares J, Kozak KH, Fine PV, Kembel SW (2009) The merging of community ecology and phylogenetic biology. Ecol Lett 12:693–715

Chase JM, Myers JA (2011) Disentangling the importance of ecological niches from stochastic processes across scales. Philos Trans R Soc Lond B Biol Sci 366:2351–2363

Da Silva MLB, Cantão ME, Mezzari MP, Ma J, Nossa CW (2014) Assessment of bacterial and archaeal community structure in swine wastewater treatment processes. Microbial Ecol 70:77–87

Dini-Andreote F, Stegen JC, van Elsas JD, Salles JF (2015) Disentangling mechanisms that mediate the balance between stochastic and deterministic processes in microbial succession. Proc Natl Acad Sci USA 112:E1326–E1332

Dridi B, Henry M, El Khechine A, Raoult D, Drancourt M (2009) High prevalence of Methanobrevibacter smithii and Methanosphaera stadtmanae detected in the human gut using an improved DNA detection protocol. PLoS ONE 4:e7063

Edgar RC, Haas BJ, Clemente JC, Quince C, Knight R (2011) UCHIME improves sensitivity and speed of chimera detection. Bioinformatics 27:2194–2200

US Environmental Protection Agency (2013) Inventory of US Greenhouse Gas Emissions and Sinks: 1990-2011. EPA Publication 430-R-13-001 (US Environmental Protection Agency, Washington, DC)

Fierer N, Jackson RB (2006) The diversity and biogeography of soil bacterial communities. Proc Natl Acad Sci USA 103:626–631

Fine PV, Kembel SW (2011) Phylogenetic community structure and phylogenetic turnover across space and edaphic gradients in western Amazonian tree communities. Ecography 34:552–565

Godon J-J, Zumstein E, Dabert P, Habouzit F, Moletta R (1997) Molecular microbial diversity of an anaerobic digestor as determined by small-subunit rDNA sequence analysis. Appl Environ Microbiol 63:2802–2813

Kembel SW, Cowan PD, Helmus MR, Cornwell WK, Morlon H, Ackerly DD et al. (2010) Picante: R tools for integrating phylogenies and ecology. Bioinformatics 26:1463–1464

Klindworth A, Pruesse E, Schweer T, Peplies J, Quast C, Horn M et al. (2012) Evaluation of general 16S ribosomal RNA gene PCR primers for classical and next-generation sequencing-based diversity studies. Nucleic Acids Res 41:e1

Kumari P, Choi HL, Sudiarto SI (2015) Assessment of bacterial community assembly patterns and processes in pig manure slurry. PLoS ONE 10:e0139437

Langenheder S, Székely AJ (2011) Species sorting and neutral processes are both important during the initial assembly of bacterial communities. ISME J 5:1086–1094

Lipp JS, Morono Y, Inagaki F, Hinrichs K-U (2008) Significant contribution of Archaea to extant biomass in marine subsurface sediments. Nature 454:991–994

Lliros M, Casamayor EO, Borrego C (2008) High archaeal richness in the water column of a freshwater sulfurous karstic lake along an interannual study. FEMS Microbiol Ecol 66:331–342

Losos JB (2008) Phylogenetic niche conservatism, phylogenetic signal and the relationship between phylogenetic relatedness and ecological similarity among species. Ecol Lett 11:995–1003

Lozupone CA, Knight R (2007) Global patterns in bacterial diversity Proc Natl Acad Sci USA 104:11436–11440

Mao SY, Yang CF, Zhu WY (2011) Phylogenetic analysis of methanogens in the pig feces. Curr Microbiol 62:1386–1389

Masella AP, Bartram AK, Truszkowski JM, Brown DG, Neufeld JD (2012) PANDAseq: paired-end assembler for illumina sequences. BMC Bioinformatics 13:31

Miller TL, Wolin M, Kusel E (1986) Isolation and characterization of methanogens from animal feces. Syst Appl Microbiol 8:234–238

Offre P, Spang A, Schleper C (2013) Archaea in biogeochemical cycles. Annu Rev Microbiol 67:437–457

Okasanen J, Blanchet FG, Kindt R, Legendre P, Minchin PR, O'Hara RB et al. (2013) vegan: Community Ecology Package. R package version 2.0-7. Available at: http://CRAN.R-project.org/package=vegan

Peay KG, Garbelotto M, Bruns TD (2010) Evidence of dispersal limitation in soil microorganisms: isolation reduces species richness on mycorrhizal tree islands. Ecology 91:3631–3640

Pei CX, Mao SY, Cheng YF, Zhu WY (2010) Diversity, abundance and novel 16S rRNA gene sequences of methanogens in rumen liquid, solid and epithelium fractions of Jinnan cattle. Animal 4:20–29

Phelps T, Conrad R, Zeikus J (1985) Sulfate-dependent interspecies H2 transfer between *Methanosarcina barkeri* and *Desulfovibrio vulgaris* during coculture metabolism of acetate or methanol. Appl Environ Microbiol 50:589–594

Price MN, Dehal PS, Arkin AP (2010) FastTree 2-approximately maximum-likelihood trees for large alignments. PLoS ONE 5:e9490

Ren L, Jeppesen E, He D, Wang J, Liboriussen L, Xing P et al. (2015) pH influences the importance of niche-related versus neutral processes in lacustrine bacterioplankton assembly. Appl Environ Microbiol 81:3104–3114

Schloss PD, Westcott SL, Ryabin T, Hall JR, Hartmann M, Hollister EB et al. (2009) Introducing mothur: open-source, platform-independent, community-supported software for describing and comparing microbial communities. Appl Environ Microbiol 75:7537–7541

Shin EC, Choi BR, Lim WJ, Hong SY, An CL, Cho KM et al. (2004) Phylogenetic analysis of archaea in three fractions of cow rumen based on the 16S rDNA sequence. Anaerobe 10:313–319

Snell-Castro R, Godon JJ, Delgenès JP, Dabert P (2005) Characterisation of the microbial diversity in a pig manure storage pit using small subunit rDNA sequence analysis. FEMS Microbiol Ecol 52:229–242

Stegen JC, Lin X, Fredrickson JK, Chen X, Kennedy DW, Murray C et al. (2013) Quantifying community assembly processes and identifying features that impose them. ISME J 7:2069–2079

Stegen JC, Lin X, Konopka AE, Fredrickson JK (2012) Stochastic and deterministic assembly processes in subsurface microbial communities. ISME J 6:1653–1664

Suresh A, Choi HL (2011) Estimation of nutrients and organic matter in Korean swine slurry using multiple regression analysis of physical and chemical properties. Bioresource Technol 102:8848–8859

Teske A, Sørensen KB (2008) Uncultured archaea in deep marine subsurface sediments: have we caught them all? ISME J 2:3–18

Torsvik V, Øvreås L, Thingstad TF (2002) Prokaryotic diversity--magnitude, dynamics, and controlling factors. Science 296:1064–1066

Tripathi BM, Kim M, Lai-Hoe A, Shukor NAA, Rahim RA, Go R et al. (2013) pH dominates variation in tropical soil archaeal diversity and community structure. FEMS Microbiol Ecol 86:303–311

Tripathi BM, Kim M, Tateno R, Kim W, Wang JJ, Lai-Hoe A et al. (2015) Soil pH and biome are both key determinants of soil archaeal community structure. Soil Biol Biochem 88:1–8

Wang JJ, Shen J, Wu YC, Tu C, Soininen J, Stegen JC et al. (2013) Phylogenetic beta diversity in bacterial assemblages across ecosystems: deterministic versus stochastic processes. ISME J 7:1310–1321

Webb CO, Ackerly DD, McPeek MA, Donoghue MJ (2002) Phylogenies and community ecology. Annu Rev Ecol Syst 33:475–505

Whitehead T, Cotta M (1999) Phylogenetic diversity of methanogenic archaea in swine waste storage pits. FEMS Microbiol Lett 179:223–226

Whitman WB, Bowen TL, Boone DR (2006) The methanogenic bacteria. Prokaryotes 3:165–207

Yan Q, Li J, Yu Y, Wang J, He Z, Van Nostrand JD et al. (2016) Environmental filtering decreases with fish development for the assembly of gut microbiota. Environ Microbiol 18:4739–4754

Zhou J, Deng Y, Zhang P, Xue K, Liang Y, Nostrand JD et al. (2014) Stochasticity, succession, and environmental perturbations in a fluidic ecosystem. Proc Natl Acad Sci 111:E836–E845

